# Neural tracking of speech does not unequivocally reflect intelligibility

**DOI:** 10.1101/2022.07.25.501422

**Authors:** Anne Kösem, Bohan Dai, James M. McQueen, Peter Hagoort

## Abstract

During listening, brain activity tracks the rhythmic structures of speech signals. Here, we directly dissociated the contribution of neural tracking in the processing of speech acoustic cues from that related to linguistic processing. We examined the neural changes associated with the comprehension of Noise-Vocoded (NV) speech using magnetoencephalography (MEG). Participants listened to NV sentences in a 3-phase training paradigm: (1) pre-training, where NV stimuli were barely comprehended, (2) training with exposure of the original clear version of speech stimulus, and (3) post-training, where the same stimuli gained intelligibility from the training phase. Using this paradigm, we tested if the neural responses of a speech signal was modulated by its intelligibility without any change in its acoustic structure. To test the influence of spectral degradation on neural tracking independently of training, participants listened to two types of NV sentences (4-band and 2-band NV speech), but were only trained to understand 4-band NV speech. Significant changes in neural tracking were observed in the delta range in relation to the acoustic degradation of speech. However, we failed to find a direct effect of intelligibility on the neural tracking of speech in both theta and delta ranges. This suggests that acoustics greatly influence the neural tracking response to speech signals, and that caution needs to be taken when choosing the control signals for speech-brain tracking analyses, considering that a slight change in acoustic parameters can have strong effects on the neural tracking response.

## Introduction

Speech presents inherent rhythmic dynamics (Ding et al., 2017; Greenberg, Carvey, Hitchcock, & Chang, 2003) to which brain activity synchronizes. Neural dynamics in the delta range (1–4 Hz) and theta range (4–8 Hz) in particular follow the slow temporal structure of speech (Ahissar et al., 2001; Gross et al., 2013; Luo & Poeppel, 2007). This neural tracking of speech is thought to be an important mechanism that would contribute to syllabic and phrasal level segmentation (Greenberg et al., 2003) therefore influencing speech perception (Giraud & Poeppel, 2012; Peelle & Davis, 2012). Yet, a longstanding debate resides on the exact mechanistic role of neural tracking in speech processing (Ding & Simon, 2014; Doelling & Assaneo, 2021; Kösem & van Wassenhove, 2017; Lakatos, Gross, & Thut, 2019; Obleser & Kayser, 2019). Speech comprehension requires a complex series of processing stages to extract meaning from sound. Therefore, neural tracking could affect speech comprehension by modulating early auditory analysis, and/or later abstract linguistic processing. In the present paper, we asked which processing stages neural tracking is involved in.

To test if neural tracking has a specific role in speech processing, experimental designs usually contrast the neural response to speech with the response to an unintelligible “control” signal. The control often results from a modulation of the clear speech’s acoustics, for instance by reversing temporally the speech signal or varying its temporal properties (Ahissar et al., 2001; Broderick, Anderson, Di Liberto, Crosse, & Lalor, 2018; Di Liberto, O’Sullivan, & Lalor, 2015a; Doelling, Arnal, Ghitza, & Poeppel, 2014; Gross et al., 2013; Howard & Poeppel, 2010; Kayser, Ince, Gross, & Kayser, 2015), by degrading the spectral resolution (Meng, Hegner, Giblin, McMahon, & Johnson, 2021; Molinaro & Lizarazu, 2018; Peelle, Gross, & Davis, 2013), or by changing the auditory background (Ding & Simon, 2013; Rimmele, Zion Golumbic, Schröger, & Poeppel, 2015; Zion Golumbic et al., 2013; Zoefel & VanRullen, 2015b). These studies mostly report that neural tracking is stronger when listening to intelligible speech as compared to unintelligible signals in both theta (Ahissar et al., 2001; Doelling et al., 2014; Peelle et al., 2013) and delta ranges (Di Liberto, O’Sullivan, & Lalor, 2015b; Ding & Simon, 2013; Doelling et al., 2014), but see (Howard & Poeppel, 2010; Zoefel & VanRullen, 2015c). However, it is still unclear from these findings whether neural tracking changes do reflect linguistic processing alone, as speech’s intelligibility covaries with acoustical changes, or whether changes in acoustics alone can modulate neural tracking (Ding, Chatterjee, & Simon, 2013; Kösem & van Wassenhove, 2017; Meng et al., 2021; Pinto, Prior, & Zion Golumbic, 2022).

In this current study, therefore, we directly dissociated the contribution of neural tracking in the processing of speech acoustic cues from those related to linguistic processing. To achieve this, we examined the neural changes associated with the comprehension of Noise-Vocoded (NV) speech (Davis, Johnsrude, Hervais-Adelman, Taylor, & McGettigan, 2005; Shannon, Zeng, Kamath, Wygonski, & Ekelid, 1995). The intelligibility of a NV sentence is dependent on its amount of spectral degradation, as directly indexed by the number of frequency bands used in the noise-vocoding procedure (Davis et al., 2005). However, the intelligibility of initially unintelligible NV speech can be recovered by training (specifically via exposure to the original clear version of the sentence) (Dahan & Mead, 2010; Sohoglu & Davis, 2016). We recorded the cortical activity using magnetoencephalography (MEG) while participants listened to NV sentences in a 3-phase training paradigm: (1) pre-training, where the NV stimulus was barely comprehended, (2) training with exposure of the original clear version of speech stimulus, and (3) post-training, where the same stimulus was more intelligible after the training phase. Using this paradigm, we tested if the neural responses to a speech signal were modulated by its intelligibility without changing its acoustic structure. To test the influence of spectral degradation on neural entrainment independently of training, participants were listening to two NV type of sentences (4-band and 2-band NV speech), but were trained to understand only 4-band NV speech.

## Materials and Methods

### Participants

Thirty-two participants were recruited. The experimental procedure was approved by the local ethics committee (CMO region Arnhem-Nijmegen), and all participants gave informed consent in accordance with the Declaration of Helsinki. All participants were right-handed native Dutch speakers, and had no known history of neurological, language, or hearing problems. One participant was excluded because she was unable to finish the experiment; leaving thirty-one participants (15 females; mean ± SD, 23 ± 3.1 years) in the analysis.

### Stimuli

We used the same NV speech stimuli as in previous behavioral and MEG studies (Dai, McQueen, Hagoort, & Kösem, 2017; Dai, McQueen, Terporten, Hagoort, & Kösem, 2022). The original speech were selected from a corpus with daily conversational Dutch sentences, digitized at a 44,100 Hz sampling rate and recorded either by a native male or a native female speaker (Versfeld, Daalder, Festen, & Houtgast, 2000). Each sentence consisted of 5-8 words (e.g., ‘Mijn handen en voeten zijn ijskoud’, in English: ‘My hands and feet are freezing’). Two semantically independent sentences recorded by the same speaker were combined into one stimulus, separated by a 300-ms silence gap (average duration = 4.15 ± 0.13 s). In total, 160 stimuli were constructed, half of them were spoken by the male speaker and half by the female speaker. The two-sentence stimuli were then manipulated by noise-vocoding (Shannon et al., 1995) with Praat software (Version: 6.0.39 from http://www.praat.org), using either 4 or 2 frequency bands logarithmically spaced between 50 and 8000 Hz, resulting in 80 trials per noise vocoding condition. The noise-vocoding technique degrades the spectral content of the acoustic signal (i.e., the fine structure) but keeps the temporal information (i.e., speech envelope) largely intact. All stimuli were normalized to ~70 dB SPL.

### Procedure

The training used in this MEG experiment was similar to our previous studies (Dai et al., 2017, 2022), but combined with more testing trials. The experiment included three phases: pre-training, training, and post-training. In the pre-training and post-training phases, the participants were tested on their ability to understand the 4-band and 2-band vocoded speech stimuli. For each trial, participants heard a speech stimulus binaurally and were asked to repeat the sentences afterwards. Participants’ responses were recorded by a digital microphone with a sampling rate of 44,100 Hz. In both pre-training and post-training phases, participants were exposed to the same 160 stimuli, with the order of presentation fully randomized in each phase. In between pre-training and posttraining, participants performed a training session to improve the intelligibility of the 4-band vocoded speech stimuli. For this, they were presented one token of the original version of the pairs of spoken sentences (i.e. the recordings on which the noise-vocoding was applied), and followed by one token of the vocoded version of that stimulus; simultaneously, to enhance the training effect, they could read the written versions of the sentences on a computer screen. 2-band vocoded speech was not trained in this phase. The experiment was implemented using Presentation software (Version 16.2, www.neurobs.com), and took about 70-80 minutes in total.

### Behavioral analysis

The intelligibility of vocoded speech was measured by calculating the percentage of correct content words (excluding function words) in participants’ reports for each trial. Words were regarded as correct if there was a perfect match (correct word without any tense errors, singular/plural form changes, or changes in sentential position). The percentage of correct content words was chosen as a more accurate measure of intelligibility based on acoustic cues than percentage correct of all words, considering that function words can be guessed based on the content words (Brouwer, 2012). A two-way repeated-measures ANOVA was performed with factors of NV band (trained 4-band and untrained 2-band) and Time (pre- and post-training). Post hoc t-tests were performed with Bonferroni correction as control for multiple comparison testing.

### MEG measurement

MEG data were recorded with a 275-channel whole-head system (CTF Systems Inc., Port Coquitlam, Canada) at a sampling rate of 1200 Hz in a magnetically shielded room. Data of four channels (MLC11, MLC32, MLF62, MRF66) were not recorded due to channel malfunctioning. Participants were seated in an upright position. Head location was measured with two coils in the ears (fixed to anatomical landmarks) and one on the nasion. To reduce head motion, a neck brace was used to stabilize the head. Head motion was monitored online throughout the experiment with a real-time head localizer and if necessary corrected between the experimental blocks. The speech signal was delivered through plastic air tubes connected to foam earpieces in the MEG scanner.

### MEG Data preprocessing

MEG Data analysis was conducted in MATLAB using the FieldTrip toolbox (fieldtrip-20190327) (Oostenveld, Fries, Maris, & Schoffelen, 2011) during pre-training and post-training sessions. Trials were defined as data between 500 ms before the onset of sound signal and 4,000 ms thereafter. Three steps were taken to remove artifacts. Firstly, trials were rejected if the range and variance of the MEG signal differed, on visual inspection, by at least an order of magnitude from the other trials of the same participant. On average, 4.2 trials (1.3%, SD = 2.3) per participant were rejected via visual inspection. Secondly, independent component analysis (ICA) was performed. Based on visual inspection of the ICA components’ time courses and scalp topographies, components showing clear signature of eye blinks, eye movement, heartbeat and noise were identified and removed from the data. Finally, 4.6 trials (1.4%, SD = 2.3) with other artifacts were removed via second visual inspection, resulting in an average of 311 included trials per participant (for both pretraining and post-training phases, namely on average 77 trials per NV band condition for each Time phase).

### MEG analysis

#### Region of Interest Analysis

A data-driven approach was first performed to identify the reactive channels for sound processing. The M100 (within the time window between 80ms and 120ms after the first word were presented) response was measured on the data over all experimental conditions, after planar gradient transformation. We selected the 6 channels with the relatively strongest response at the group level on each hemisphere, and the averages of these channels were used for all subsequent analysis.

Speech-brain coherence analysis was performed on the data within the region of interest after planar gradient projection. Spectral analysis was performed using ‘dpss’ multi-tapers with a ± 1 Hz smoothing window of the speech envelopes, and of the neural times series epoched from 500 epochs were removed to exclude the evoked response to the onset of the sentence. To match trials in duration, shorter trials were zero-padded up to 4s. The speech-brain coherence was measured at different frequencies (1 to 30 Hz, 1 Hz step). Finally, the coherence data were projected into planar gradient representations. Then data were averaged across all trials and the strongest 12 channels defined by our auditory response localizer. For the investigation of our main hypotheses, we restricted the speech-brain coherence analyses to delta band (1–4 Hz) and theta band (4–8 Hz) activity; these frequency bands were chosen based on the previous literature (Ding & Simon, 2014; Kösem & van Wassenhove, 2017). We repeated the same analysis described above to quantify the speech-brain coherence for each condition, and the averaged speech-brain coherences of the strongest 6 channels on each hemisphere in the delta and theta bands were calculated. We tested the speech-brain coherence in the delta and theta range using a two-way repeated measure ANOVA with factors NV band (4-band, 2-band), Time (pre-training, post-training).

#### Whole sensor space analysis

We performed cluster-based permutation statistics across subjects (Oostenveld et al., 2011) to test whether we could observe a main effect of NV-band across sensors on speech-brain coherence *Coh* (by contrasting between *Coh_4-band_* and *Coh_2-band_*, averaged across pre- and post-training sessions) and an interaction effect between NV-band and Time (by contrasting between (*Coh_4-band, post_ - Coh_4-band, pre_*) and (*Coh_2-band, post_ - Coh_2-band, pre_*) in both delta and theta frequency ranges. Pairwise t-tests were then computed for each sensor between the two conditions. Sensors with a p-value associated to the t-test of 5% or lower were selected as cluster candidates. The sum of the t-values within a cluster was used as the cluster-level statistic. The reference distribution for cluster-level statistics was computed by performing 1,000 permutations between the two conditions. Clusters were considered significant if the probability of observing a cluster test statistic of that size in the reference distribution was 0.025 or lower (two-tailed test).

#### Source reconstruction analysis

Anatomical MRI scans were obtained after the MEG session using either a 1.5 T Siemens Magnetom Avanto system or a 3 T Siemens Skyra system for each participant (except for two participants whose data were excluded from source reconstruction analysis). The co-registration of MEG data with the individual anatomical MRI was performed via the realignment of the fiducial points (nasion, left and right pre-auricular points). Lead fields were constructed using a single shell head model based on the individual anatomical MRI. Each brain volume was divided into grid points of 1 cm voxel resolution, and warped to a template MNI brain. For each grid point the lead field matrix was calculated. The sources of the observed delta and theta speech-brain coherence were computed using beamforming analysis with the dynamic imaging of coherent sources (DICS) technique to the coherence data.

## Results

### Behavioral results

We compared the participants’ comprehension of NV speech before and after training. Consistent with previous findings (Dai et al., 2017, 2022; Davis et al., 2005; Sohoglu & Davis, 2016), and as shown in Fig. 2, the training significantly improved the perception of 4-band NV speech. A twoway repeated-measure ANOVA showed that the main effects of noise vocoding (2-band *vs*. 4-band) and time (pre- *vs*. post-training) were significant (noise vocoding: (*F*(1, 30) = 331.48, *p* < 0.001, *η^2^* = .92; time: *F*(1,30)= 183.46, *p* < 0.001, *η^2^* = .86). Crucially, a significant interaction between noise vocoding and time was observed (*F*(1,30) = 180.99, *p* < 0.001, *η^2^* = .86), meaning that the intelligibility of 4-band NV speech was significantly improved compared to that of 2-band NV speech (4-band(post-pre) vs. 2-band(post-pre): *t* (24) = 13.45, *p* < 0.001, Cohen’s *d* = 2.42). After training, 4-band NV sentences had a score of 43.38 ± 2.53% recognition accuracy (23.06 ± 1.69% improvement during training; values here and below indicate mean ± SEM), while 2-band NV sentences remained mostly unintelligible with a score of 2.39 ± 0.43% recognition accuracy (1.58 ± 0.28% improvement during training).

**Figure 1.**
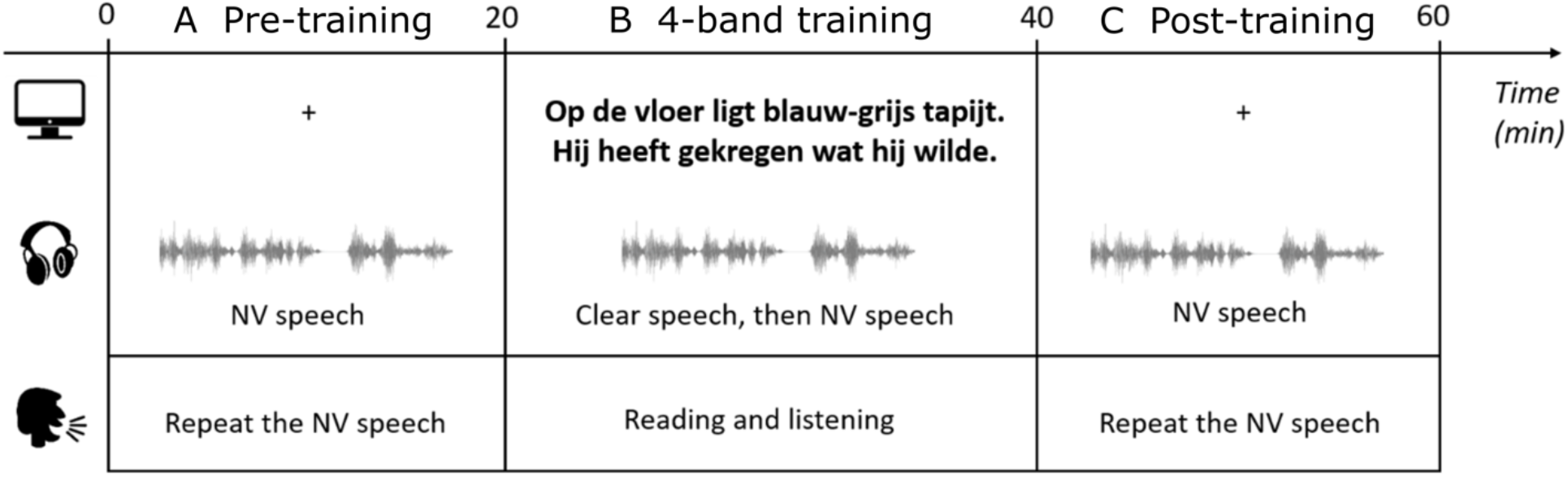
Experimental design. The experiment consisted of three phases: pre-training (A), 4-band training (B), and post-training (C). In the pre- and post-training phases, the participants were tested on their ability to understand the 4-band and 2-band vocoded speech stimuli. They were presented with the speech signal binaurally and were asked to report the sentences afterwards. During the training phase, participants listened to clear-speech versions of the 4-band pre-training sentences followed by the NV versions. At the same time, they read the text of the sentences on the screen.

**Figure 2:**
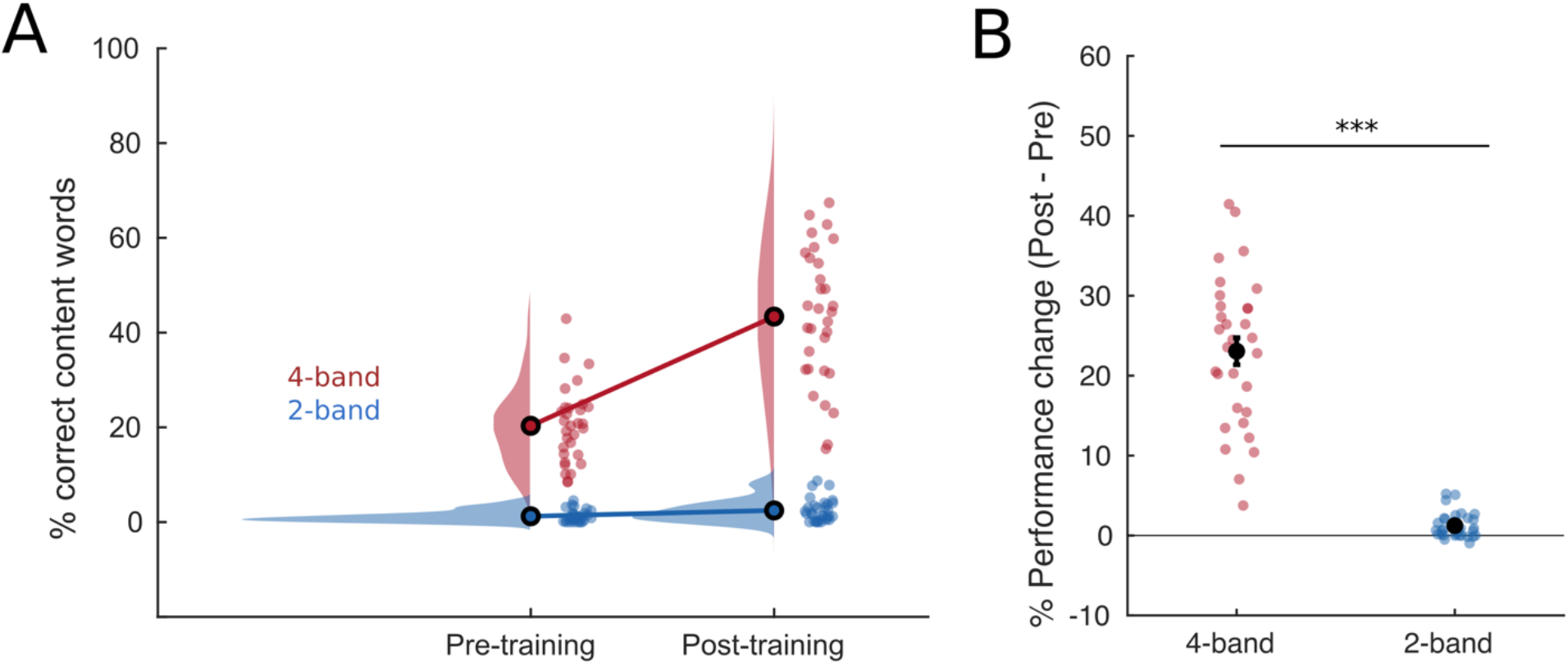
Behavioral results. (A) Proportion of corrected reported content words pre- and post-training for both NV-speech conditions. (B) Performance change Post – Pre training. The intelligibility of trained 4-band (red) significantly improved by 23% on average with training, while untrained 2-band (blue) NV speech remained mostly unintelligible post-training. The open dots connected by lines (panel A) and the large black dots (panel B) indicate the grand average performance in each condition. The rainclouds indicated the distributions of individual data, and each small dot corresponds to one participant.

### MEG results

The behavioral results confirmed that intelligibility and spectral complexity could be dissociated in the present study. We then investigated how speech-brain coherence in auditory regions was impacted by the training session (Fig. 3). In line with previous studies (Meng et al., 2021; Peelle et al., 2013), we show that the neural tracking of 4-band NV speech was stronger than that to 2-band NV speech (Fig. 3B). This was observed in the delta but not the theta frequency range (main effect of NV band, delta: F(1, 30) = 25.29, p < .001, η^2^ = .46; theta: F(1, 30) = 1.43, p = .24, η^2^ = .04, Fig 3C-D).

**Figure 3:**
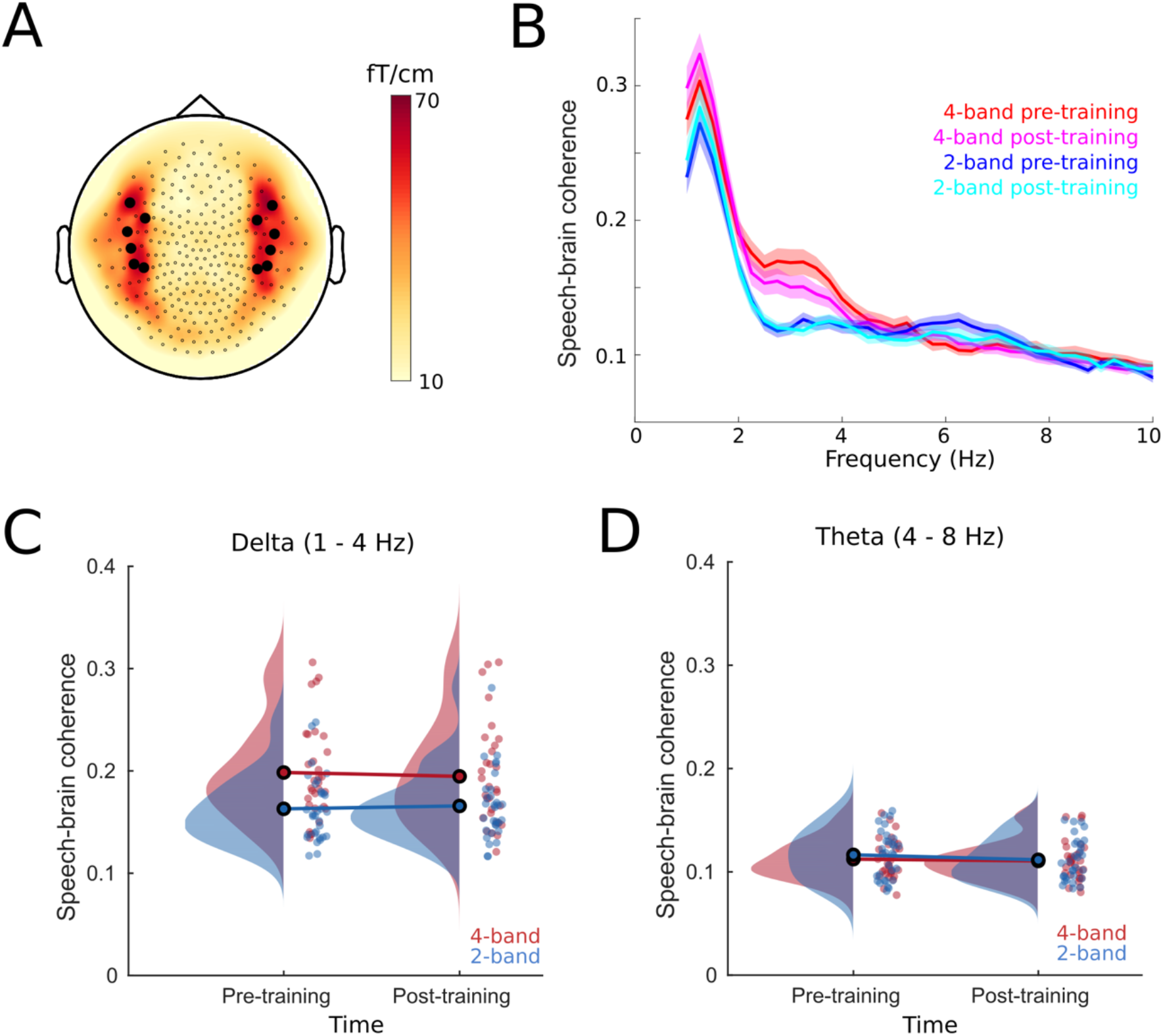
Neural tracking responses in auditory cortices as a function of intelligibility and acoustic spectral complexity. (A) Topography of the M100 response. The highlighted six channels showed the relatively strongest response at the group level on each hemisphere, and the average within these channels was used for all subsequent region-of-interest analysis. (B) Speech-brain coherence averaged across selected channels. (C) Coherence between neural activity and 4-band NV speech (red) or 2-band NV speech (blue) in the delta (1-4 Hz) range averaged across selected channels. The open dots connected by lines indicate the grand average speech brain coherence in the pre- and post-training phases. The rainclouds indicated the distribution of individual data, and each small dot corresponds to one participant. (D) Coherence between neural activity and 4-band NV speech (red) or 2-band NV speech (blue) in the theta (4-8 Hz) range averaged across selected channels.

However, the neural tracking of NV speech was not significantly affected by training at either delta or theta frequencies (Fig. 3C, delta, main effect of time: (F(1, 30) = .03, p = .85, η^2^ = .001; Fig. 2D & F, theta: (F(1, 30) = 2.87, p = .10, η^2^ = .09)). If neural entrainment reflected intelligibility, we specifically predicted that neural entrainment to NV speech signals would be stronger after training for the intelligible 4-band NV sentences. Yet, this was not observed (interaction between NV-band and Time, delta: F(1, 30) = 2.43, p = .13, η^2^ = .07; theta: F(1, 30) = 0.53, p = .47, η^2^ = .02).

Further whole brain analysis showed a similar pattern of results (Fig. 4A-B). Cluster-based permutation tests revealed a main effect of NV-band in the delta range (cluster p <0.001), but not in the theta range. No significant interaction effects were observed. Overall, the results suggest that speech-brain coherence is influenced by the acoustic structure of the speech signal, rather than reflecting linguistic processing.

**Figure 4:**
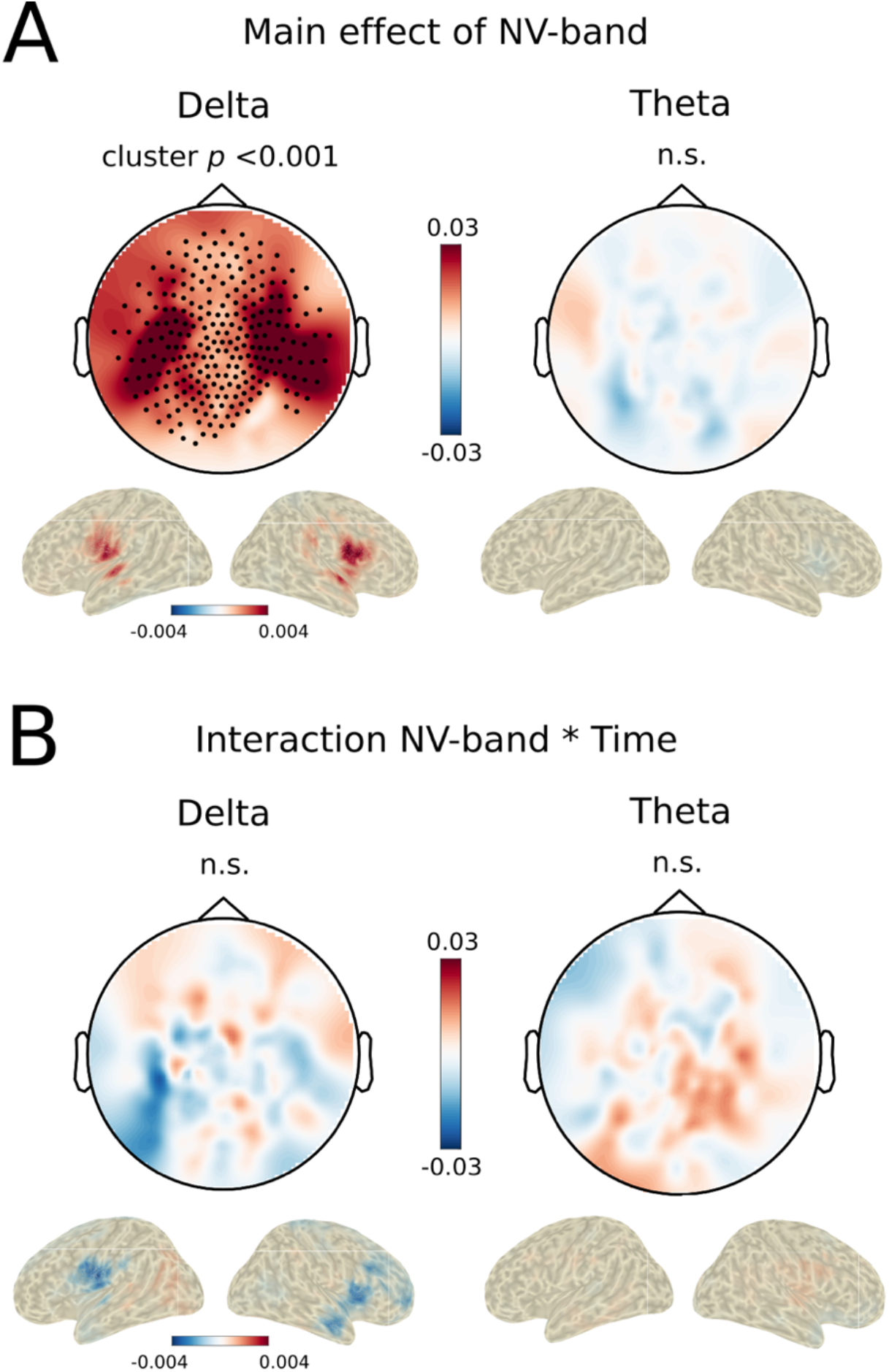
Whole brain analysis. (A) Main effect of NV-band. Topographies and reconstructed source represent the contrast in neural tracking of 4-band vs 2-band NV speech. Left panel: A main effect of NV-band is observed in the delta range (dots represent spatial topography of the significant cluster). Delta tracking is stronger for 4-band NV speech as compared to 2-band NV, irrespective of training. Right panel: No significant changes in neural tracking of speech in the theta range irrespective of their spectral complexity. (B) Interaction effects NV-band * Time. No significant effect of training is observed, specifically the gain in intelligibility of the 4-band NV speech after training is not associated with a gain in neural tracking.

## Discussion

In the present study, we tested the effect of intelligibility on the neural tracking of speech. We used NV speech that could gain in intelligibility via training. With this manipulation we could dissociate gains in intelligibility linked to acoustic cues (spectral degradation), from those linked to linguistic processing of the speech signal. The training increased the intelligibility of NV speech but did not change its neural tracking response. In contrast, neural tracking in the delta range was still modulated by the acoustic detail of the NV speech signal. These results are in line with previous reports showing that neural tracking reduces with the amount of spectral degradation (Meng et al., 2021; Peelle et al., 2013), and others failing to find a correlation between the neural tracking of speech in auditory cortex and speech intelligibility when acoustic details in speech are controlled (Kösem, Basirat, Azizi, & van Wassenhove, 2016; Millman, Johnson, & Prendergast, 2015; Peña & Melloni, 2012; Zoefel & VanRullen, 2015a). Therefore, the results suggest that brain-speech tracking reflects relevant neural mechanisms during the processing of speech acoustics, but does not unequivocally reflect the processing of more abstract linguistic information in speech.

This interpretation seems in apparent contradiction with other findings. Ding and colleagues (Ding et al, 2016) have found that neural oscillations in the delta range could track sentential and phrasal linguistic structures in speech in the absence of acoustic cues. A recent study reanalyzing the data of (Millman et al., 2015) has found that delta tracking of speech is increased when the NV speech is intelligible as compared to when it is not understood (Di Liberto, Lalor, & Millman, 2018). In noisy environments, neural tracking of the attended speech signal is stronger when the attended speech is fully understood (Dai et al., 2022; Keitel, Gross, & Kayser, 2018), or when the attended speech is in competition with unstructured speech (words were presented in random order) as compared to structured speech (speech with phrasal structure) (Har-Shai Yahav & Zion-Golumbic, 2021). The language proficiency of the listener also affects the neural tracking of naturally spoken speech (Lizarazu, Carreiras, Bourguignon, Zarraga, & Molinaro, 2021).

One difference between these other studies and the present one concerns the intelligibility level of the stimuli. In the prior studies, intelligibility ratings were very high as compared to our design, where maximum intelligibility reached 40-60%. This means that our participants may have learned to extract some phonological and lexical cues from the speech, but may not have enough information to extract the full content of the sentences or be able to predict their linguistic structure. In contrast, in the prior studies, the intelligible stimuli were understood for the most part. Moreover, the syntactic structure of the stimuli was clearly predictable in some experimental designs: in Ding, Melloni, Zhang, Tian, & Poeppel (2016) sentences with similar phrasal and sentential structure were presented in blocks; in Di Liberto et al. (2018) the same sentence was repeated over and over. Therefore, delta tracking may reflect the processing of intelligible and predictable linguistic information (as in the prior studies), but may not do so (as in the current study) when the speech signal is too noisy, does not have a predictable syntactic structure, and/or is not fully intelligible.

It is also important to point out that, in the prior studies mentioned above, the effect of intelligibility on brain-speech tracking seemed to be restricted to delta dynamics (< 4 Hz) and was less clearly observable for theta dynamics. These data supports the predominant role in delta tracking in the processing of linguistic structure, while theta tracking may affect the processing of acoustic and phonological information (Kösem & van Wassenhove, 2017). Still, we did not find an effect of intelligibility in delta dynamics, and we show that spectral degradation differently affected delta and theta neural tracking of speech. The increased spectral degradation of speech was associated with decreased delta tracking in auditory areas, while theta tracking remained unaffected by the amount of noise vocoding. These results suggest that theta dynamics may primarily track broadband envelope temporal information (that is unaffected by the amount of vocoding), while neural tracking of speech in the delta range may be impacted by the spectral complexity of the speech signal (Ding et al., 2013; Meng et al., 2021).

In conclusion, we failed to find a direct effect of intelligibility on the neural tracking of speech in both theta and delta ranges. Significant changes in neural tracking were still observed in the delta range in relation to the acoustic degradation of speech. These findings suggest that acoustics greatly influence the neural tracking of speech signals. They also suggest that caution is required when choosing the control condition for analyses of tracking responses because, as we have shown, a slight change in acoustic parameters can have strong effects on the neural tracking response. Finally, they suggest that neural tracking is not necessarily modulated by the intelligibility of the speech signal.

## Acknowledgments

This research was supported by a Spinoza award to P.H. and by a Marie Skłodowska-Curie Individual Fellowship (grant 843088) to A.K.

